# FTO promotes skeletal muscle differentiation and regeneration by regulating m^6^A-modified c-Myc

**DOI:** 10.64898/2025.12.30.696954

**Authors:** Paromita Dey, Bijan K. Dey

**Affiliations:** The RNA Institute, University at Albany, State University of New York (SUNY), 1400 Washington Avenue, Albany, New York 12222; Department of Biological Sciences, University at Albany, State University of New York (SUNY), 1400 Washington Avenue, Albany, New York 12222

**Keywords:** FTO, c-Myc, m^6^A, muscle stem cells, myoblast, skeletal muscle, differentiation, regeneration

## Abstract

Recent studies have revealed the crucial role of m^6^A RNA methylation in various myogenic processes. However, the specific function and underlying molecular mechanisms of this modification *in vivo* during skeletal muscle differentiation and regeneration remain unclear. In this study, we examine the role and mechanism of the m^6^A RNA demethylase fat mass and obesity-associated protein (FTO) in skeletal muscle differentiation and regeneration in mice. Our findings demonstrate that FTO is upregulated during both skeletal muscle differentiation and regeneration and is essential for these key myogenic processes. We show that exogenous FTO expression in primary myoblasts enhances differentiation, whereas FTO knockdown inhibits it. Additionally, FTO knockout in mouse muscle stem cells impairs muscle regeneration. FTO promotes skeletal muscle differentiation and regeneration by directly targeting and regulating m^6^A-modified c-Myc, a well-known repressor of myogenesis. The IGF2BP2 reader protein recognizes m^6^A-modified c-Myc in undifferentiated myoblasts and stabilizes it. As FTO levels increase during myoblast differentiation, m^6^A levels on c-Myc decrease. This reduction prevents IGF2BP2 from binding to c-Myc, thereby destabilizing c-Myc levels and promoting differentiation. Overall, our findings underscore the significance of the novel FTO/c-Myc/IGF2BP2 axis in skeletal muscle differentiation and regeneration.

## Introduction

Skeletal muscle stem cells (MuSCs), located on muscle fibers, are usually in a state of mitotic quiescence^1^. However, during skeletal muscle development and regeneration after birth, quiescent MuSCs become activated and re-enter the cell cycle^1^. Once activated, they proliferate to generate a pool of myoblasts. These myoblasts then differentiate and fuse to form new myofibers or repair existing ones. Some activated MuSCs can self-renew, returning to a quiescent state to maintain the MuSC pool^1^. This complex process, known as myogenesis, is vital for maintaining normal physiological function. Myogenesis depends on tightly regulated gene expression to ensure proper skeletal muscle function. Disruptions in gene expression during critical cellular events in skeletal myogenesis can lead to severe muscle degenerative diseases.

Much of the current understanding of key myogenic processes, including myoblast differentiation and skeletal muscle regeneration, stems from studies of the regulation of myogenic transcription factors and signaling molecules. For example, the PAX7 transcription factor is expressed and necessary for MuSC survival, proliferation, and the prevention of premature differentiation^2,3^. Loss of PAX7 and its homologue, PAX3, impairs MuSC survival and prevents MuSCs from entering the non-myogenic lineage^4^. PAX3 and PAX7 act upstream of MYOD to activate myogenic gene expression and initiate the myogenic program^2^. MYOD and MYF5 are upregulated during MuSC activation. MYOG and MRF4 are upregulated during early myogenesis, and MHC is upregulated during late myogenic differentiation. Studies in mice show that MYF5 and MYOD are essential for myogenic commitment, MYOG is required for myotube formation, and MRF4 is required for myofiber maturation^5^. Key signaling molecules, such as BMPs, Wnts, AKT, and mTOR, are vital for normal myogenesis^6,7^. However, the gene regulatory mechanisms that control these transcription factors and signaling molecules within the myogenic network, which are crucial for normal muscle formation and function, remain poorly understood. Therefore, identifying novel gene regulatory mechanisms that control these critical myogenic factors is vital to advancing our understanding of myogenesis.

In this context, there is growing interest in the role of epitranscriptomics, a new mode of post-transcriptional gene regulation during skeletal myogenesis. Epitranscriptomics examines how enzyme-mediated chemical modifications on messenger RNAs (mRNAs) influence a wide range of biological functions. N^6^-methyladenosine (m^6^A) is the most abundant mRNA modification and is conserved across species^8,9^. The formation of m^6^A is catalyzed by a multi-protein complex that includes methyltransferase-like 3 (METTL3)^8^and methyltransferase-like 14 (METTL14)^8^. METTL3 possesses the catalytic activity^10^, while METTL14 recognizes the substrate^11,12^. Additionally, WTAP, VIRMA, RBM15/15B, ZC3H13, HAKAI, and KIAA1429 are key proteins associated with the METTL3-METTL14 complex^13–15^. The m^6^A epitranscriptomic marks are erased by fat mass and obesity-associated (FTO)^16^ and alkylation repair homolog 5 (ALKBH5)^17^. The YTH domain-containing reader proteins (YTHDF1-3, YTHDC1-2) and IGF2 mRNA-binding proteins 1, 2, and 3 (IGF2BP1-3) selectively recognize m^6^A on mRNAs and regulate their fate, including degradation or stabilization of transcripts and translation^15,18–23^.

The function of m^6^A RNA methylation has recently been linked to myogenesis^24–33^. METTL3 has been shown to regulate MuSC/myoblast state transitions^25^, maintain *Myod* mRNA in proliferating myoblasts^26^, influence myoblast differentiation^25,27,28^, regeneration^27^, skeletal muscle maintenance and growth^29^, and regulate fat intake in skeletal muscles^30^. ALKBH5 has been demonstrated to promote denervation-induced skeletal muscle atrophy^33^. Although the detailed molecular mechanisms of m^6^A RNA methylation in myogenesis remain poorly understood, m^6^A influences these essential processes by modifying and regulating key myogenic factors and signaling molecules via reader proteins^28,34^.

As the name suggests, FTO was initially linked to obesity-related traits and risk through single-nucleotide polymorphisms (SNPs) within it^35^. Loss of FTO results in reduced body weight and lean mass, as well as increased obesity^36,37^. Subsequently, FTO was identified as a demethylase involved in m^6^A RNA methylation^24^. FTO-dependent demethylation of m^6^A contributes to fat accumulation in skeletal muscle^38^, muscle fiber remodeling^39^, and myoblast differentiation *in vitro*^24,32^. However, the role of FTO and its underlying molecular mechanisms in skeletal muscle differentiation and regeneration *in vivo* remains unknown. In this study, we reveal the pivotal role and intricate mechanism of FTO in mouse skeletal muscle differentiation and regeneration. Our findings demonstrate that FTO is significantly upregulated during both the differentiation and regeneration phases of skeletal muscle, underscoring its essential role in these critical myogenic processes. Exogenous expression of FTO in myoblasts enhances differentiation, whereas inhibition of FTO suppresses it. Similarly, conditional knockout of FTO in mouse MuSCs impairs muscle regeneration. FTO promotes skeletal muscle differentiation and regeneration by directly targeting and regulating m^6^A-modified *c-Myc*, a repressor of myogenesis^40^. This process is facilitated by the IGF2BP2 reader protein, which recognizes and stabilizes m^6^A-modified *c-Myc* in undifferentiated myoblasts. As FTO expression increases during myoblast differentiation, m^6^A modifications on *c-Myc* decrease, preventing IGF2BP2 from stabilizing *c-Myc* and thereby promoting differentiation. These findings underscore the crucial role of the novel FTO/c-Myc/IGF2BP2 axis in regulating skeletal muscle differentiation and regeneration.

## Results

### FTO is highly expressed in skeletal muscles and is upregulated during myoblast differentiation and skeletal muscle regeneration

To establish a link between FTO and myogenesis, we first examined FTO expression in skeletal muscle and compared it with levels in various mouse tissues. Our results showed that FTO is highly expressed in adult skeletal muscle compared with other tissues (**Fig. 1A**). Next, we analyzed RNA-seq datasets from undifferentiated and differentiated C2C12 myoblasts and found that FTO expression is upregulated during myoblast differentiation (**Suppl. Fig. 1A**). We confirmed these RNA-seq findings using quantitative reverse transcriptase polymerase chain reaction (qRT-PCR) and Western blot analyses of FTO. We observed that FTO expression was indeed elevated at both the transcript and protein levels during C2C12 myoblast differentiation (**Suppl. Fig. 1B, C**). We then extended these findings to mouse primary myoblasts, which are physiologically more relevant for studying myogenesis. Similar to C2C12 cells, FTO expression was also upregulated at both the transcript and protein levels during primary myoblast differentiation (**Fig. 1B, C**).

**Figure 1:**
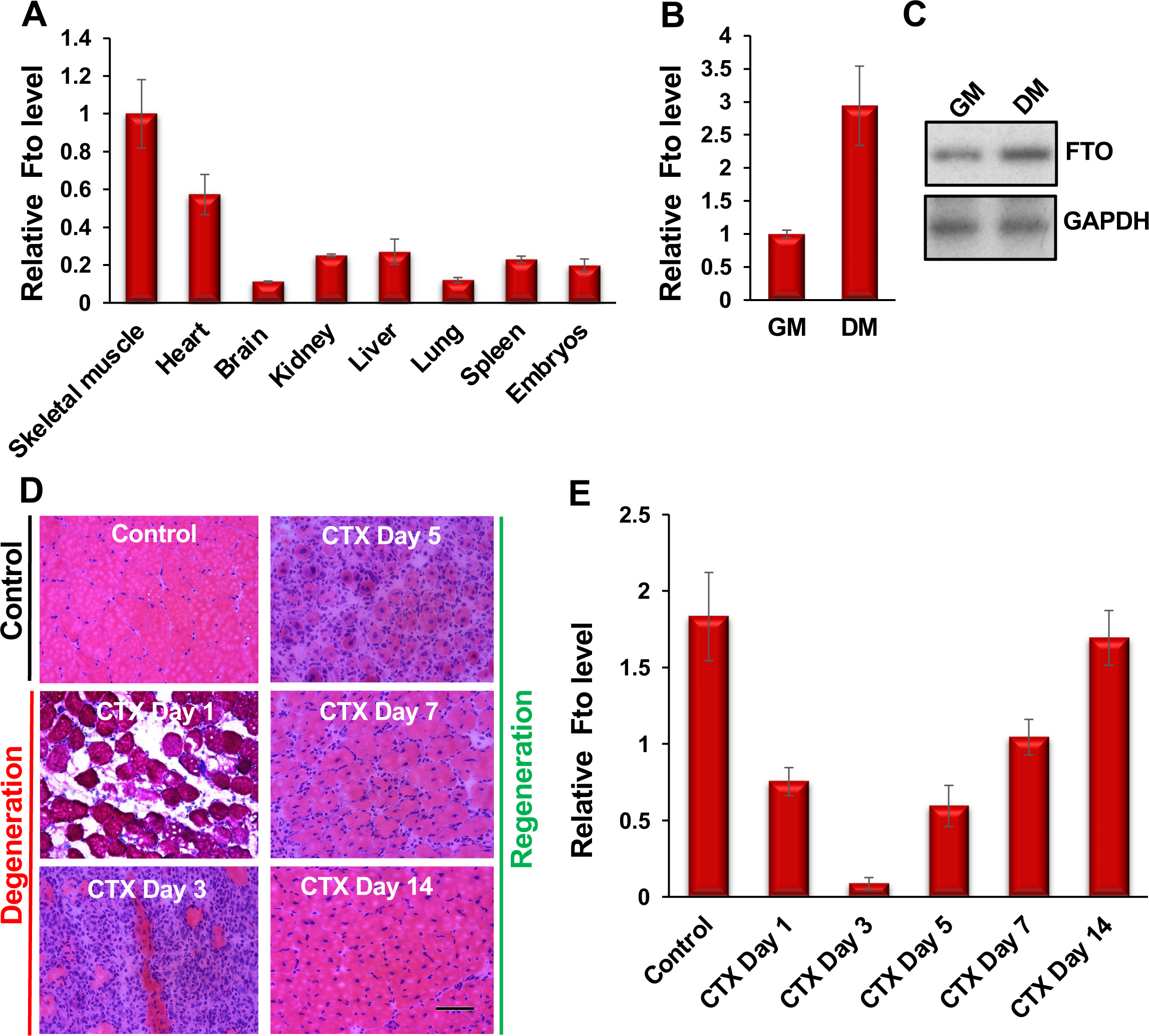
FTO is highly expressed in adult skeletal muscles and is regulated during myoblast differentiation and muscle regeneration. (**A**) *Fto* is abundantly expressed in adult skeletal muscles. *Fto* levels were measured in various mouse tissues by qRT-PCR. *Fto* values were normalized to the respective *Gapdh* values and then adjusted relative to skeletal muscle values. (**B**) qRT-PCR analyses show that *Fto* is upregulated during primary myoblast differentiation. The *Fto* level was first normalized to *Gapdh*, and fold changes were calculated relative to undifferentiated primary myoblasts (GM). DM indicates primary myoblasts cultured in differentiation medium (DM) for 72 hours. (**C**) FTO protein level, as shown by Western blotting, is also increased during mouse primary myoblast differentiation. GAPDH served as a loading control. (**D**) Hematoxylin and eosin (H&E) staining of tibialis anterior (TA) muscle cross-sections from noninjected controls and at days 1, 3, 5, 7, and 14 post-injury caused by intramuscular injection of cardiotoxin (CTX). (**E**) *Fto* is downregulated on days 1–3 post-injury (the degeneration phase) and upregulated on days 5–14 post-injury (the regeneration phase). The values are shown as mean ± SD of three (A, B) and five (E) biological replicates.

To explore the role of FTO in muscle regeneration, we examined FTO expression during skeletal muscle degeneration and regeneration using a well-established mouse model. We injected cardiotoxin (CTX) into the tibialis anterior (TA) muscles of 10-week-old adult mice and monitored the animals at multiple time points. After a CTX injury, mature muscle fibers undergo degeneration. During this phase, quiescent MuSCs are activated, giving rise to myoblasts. This is followed by a regeneration phase, in which myoblasts differentiate and fuse with existing myofibers to repair the injured myofiber or form new myofibers (**Fig. 1D**). FTO levels decrease rapidly during days 1 to 3 (the degeneration phase) and then increase from days 5 to 14 (the regeneration phase) (**Fig. 1E**). These findings suggest that FTO is regulated during myoblast differentiation and skeletal muscle regeneration, indicating its role in myogenesis.

### FTO promotes myoblast differentiation and is crucial for facilitating this process

Myoblast differentiation is a crucial step in postnatal muscle development and regeneration. Therefore, we investigated the role and requirements of FTO in primary myoblast differentiation using both exogenous FTO expression and FTO knockdown. We expressed FTO exogenously using a retroviral system following our established protocol. Briefly, primary myoblasts were transduced overnight with viral particles encoding the FTO expression vector. The next day, the medium was replaced with selection medium containing antibiotics to select transduced cells. This was replaced after 36 hours. The selected transduced cells were plated on coverslips in DM medium and harvested after an additional 24 hours for MYOG immunocytochemistry and 48 hours for MHC immunocytochemistry. We observed that increased FTO expression in these cells (**Suppl. Fig. 2A**) was associated with higher numbers of MYOG- and MHC-positive cells (**Fig. 2A-C, Suppl. Table 1**). Additionally, increased FTO levels in these cells elevated *Myog* and *Mhc* transcripts, even when the cells were maintained in GM throughout (**Fig. 2D**). MYOG and MHC protein levels also increased in these cells (**Fig. 2E**). These findings suggest that FTO can promote myoblast differentiation.

**Figure 2:**
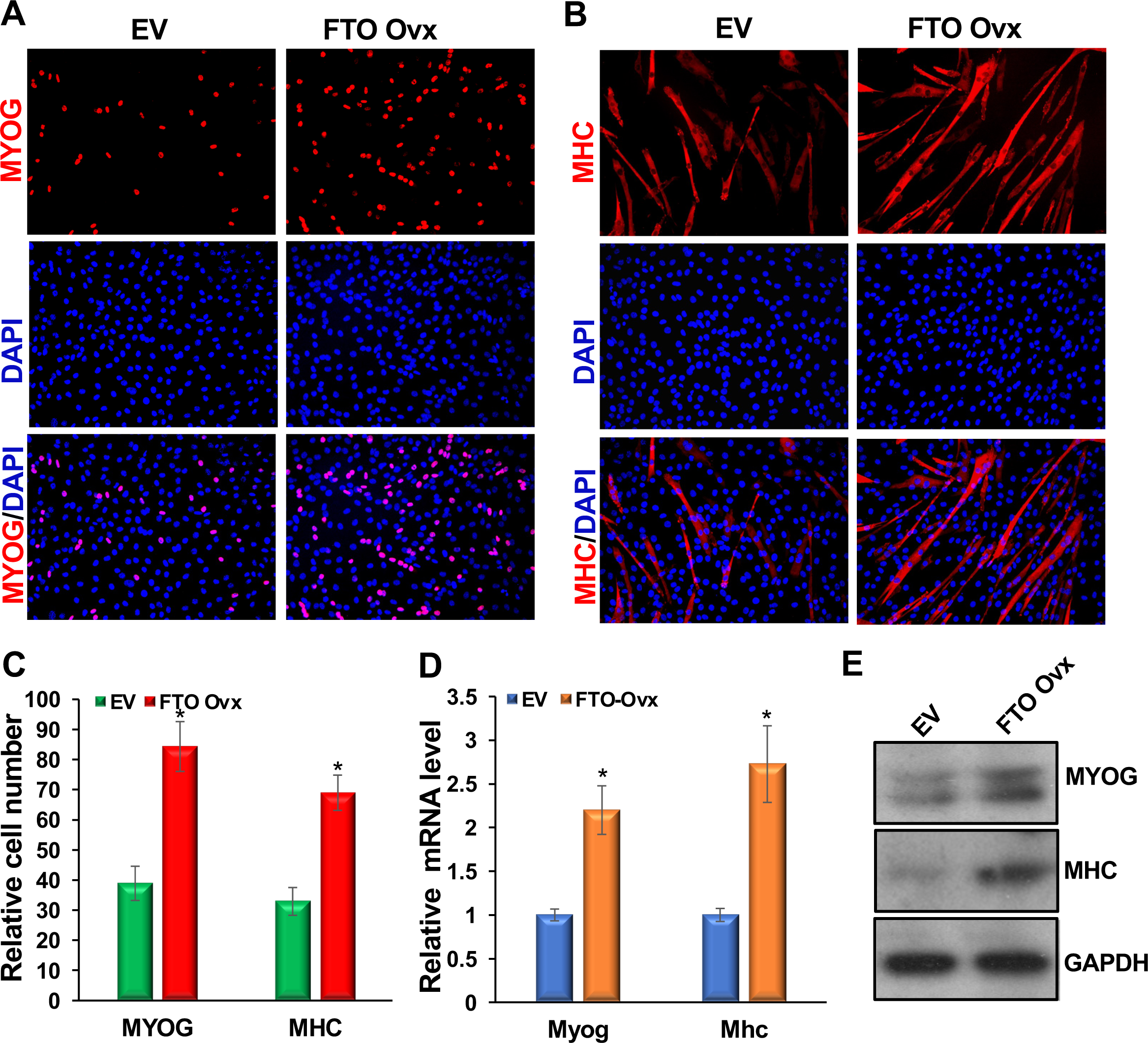
FTO promotes mouse primary myoblast differentiation. Mouse primary myoblasts were transduced with retroviral particles containing the FTO expression vector, and stable cells were selected. The medium was then replaced with DM. (**A, B**) Immunostaining for MYOG and MHC was performed on these cells at 24 and 48 hours after medium replacement, respectively. (**C**) Relative MYOG- and MHC-positive cell counts are shown. (**D**) *Myog* and *Mhc* mRNA levels were measured while cells were maintained in GM throughout. qRT-PCR for *Myog* and *Mhc* was performed, and results were normalized to *Gapdh* and then to empty vector (EV)-transduced negative control values. (**E**) MYOG and MHC protein levels. GAPDH served as a loading control. Values are expressed as mean ± SD of biological triplicates. *P < 0.001. For MYOG- and MHC-positive cells, four random fields from each biological sample were counted, for a total of 12 fields.

Next, we examined the requirement for FTO in myoblast differentiation using knockdown experiments. We used Locked Nucleic Acid (LNA) GapmeR inhibitors specific to FTO to knock down FTO levels. LNA GapmeRs are chemically modified small RNA oligonucleotides that target and degrade specific RNA sequences via the RNase H pathway. They offer several advantages, including increased stability, reduced off-target effects due to higher affinity for the target RNA, and efficient uptake by animal cells, resulting in more effective gene silencing. First, we incubated myoblasts with either FTO-specific GapmeRs (gap-FTO) or negative-control GapmeRs (gap-NC) twice at 24-hour intervals in growth medium (GM). After 24 hours, we replaced the GM with differentiation medium (DM). We then performed immunostaining for MYOG and MHC at 24 and 48 hours after DM addition. Our results showed that gap-FTO significantly reduced FTO levels (**Suppl. Fig. 2B**), decreased the number of MYOG- and MHC-positive cells, and altered differentiation morphology (**Fig. 3A-C; Suppl. Table 2**). Additionally, we observed that *Myog* and *Mhc* transcript and protein levels were substantially decreased in these samples (**Fig. 3D, E**). These findings demonstrate that FTO is essential for normal myoblast differentiation.

**Figure 3:**
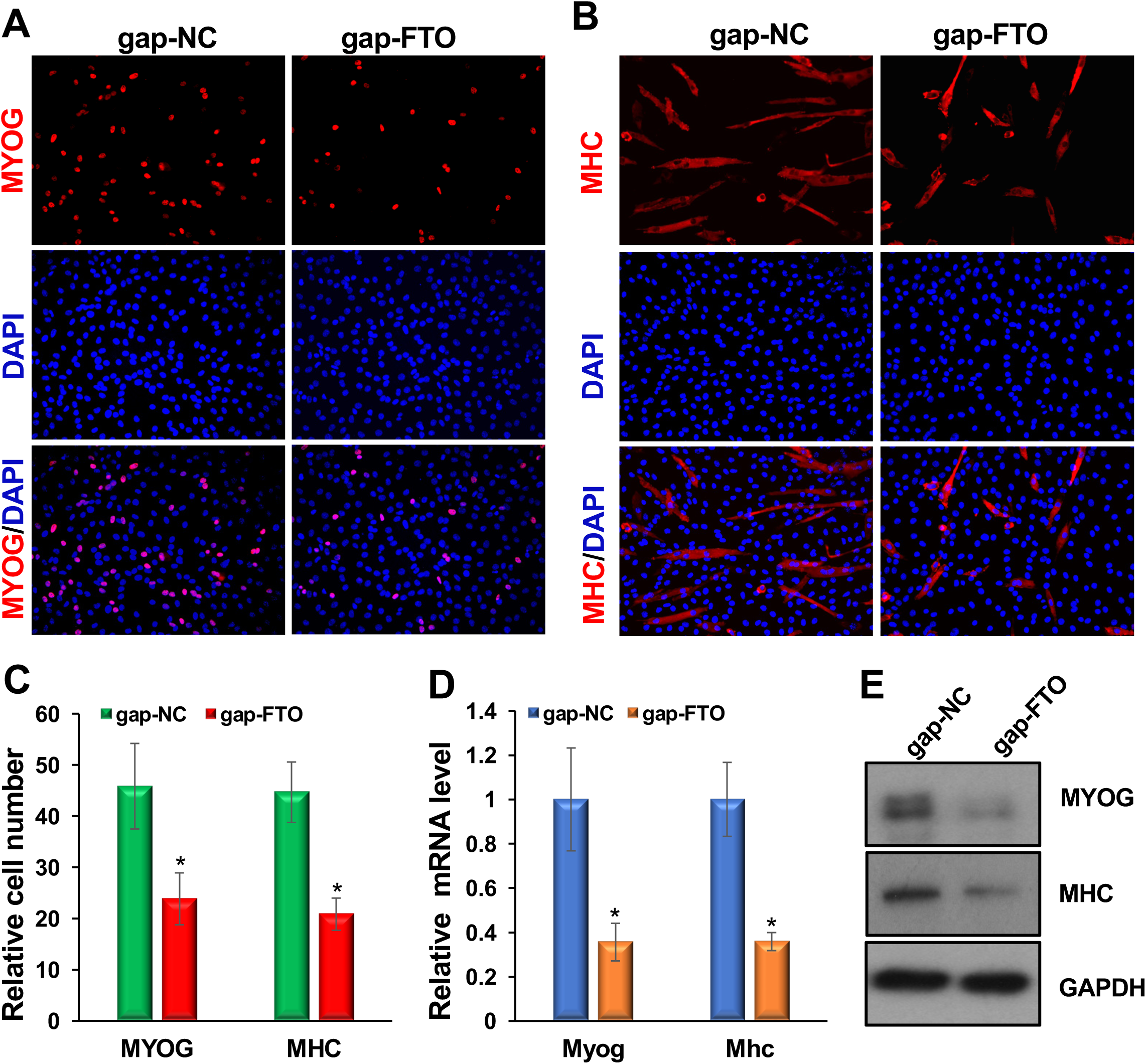
FTO is essential for mouse primary myoblast differentiation. Primary myoblasts were transfected twice at 24-hour intervals with either gap-FTO or gap-NC, and the GM was then replaced with DM. (**A, B**) Immunostaining for MYOG and MHC was performed 24 and 48 hours after medium replacement, respectively. The gap-FTO significantly reduced the number of MYOG- and MHC-positive cells and altered their differentiation morphology. (**C**) Relative numbers of MYOG- and MHC-positive cells are shown. (**D**) qRT-PCR for *Myog* and *Mhc* was performed, and results were normalized to *Gapdh* and then to gap-NC values. (**E**) Protein levels of MYOG and MHC are shown. GAPDH served as a loading control. Values are expressed as mean ± SD of biological triplicates. *P < 0.001. For MYOG- and MHC-positive cells, four random fields from each sample were counted, for a total of 12 fields.

### FTO promotes myogenesis by regulating m^6^A-modified c-Myc

Next, we investigated how FTO promotes myogenesis. To address this mechanistic question, we sought to identify FTO-regulated m^6^A-modified RNAs that play a critical role in myogenesis. We screened a set of transcription factors and signaling molecules involved in myogenesis using the SRAMP bioinformatics algorithm to identify potential m^6^A-modified RNAs^41^. We identified *c-Myc* as a potential m^6^A-modified target RNA (**Fig. 4A**). The SRAMP algorithm detected a cluster of very high-confidence m^6^A sites near the stop codon of the *c-Myc* transcript, consistent with the m^6^A consensus DRACH sequence. We chose *c-Myc* because of its essential functions in myogenesis to further validate whether it is m^6^A-modified and regulated by FTO. *c-Myc* is highly expressed in proliferating myoblasts and is downregulated during myoblast differentiation. It has been shown to inhibit myoblast differentiation by repressing promyogenic microRNAs and long noncoding RNAs^42^.

**Figure 4:**
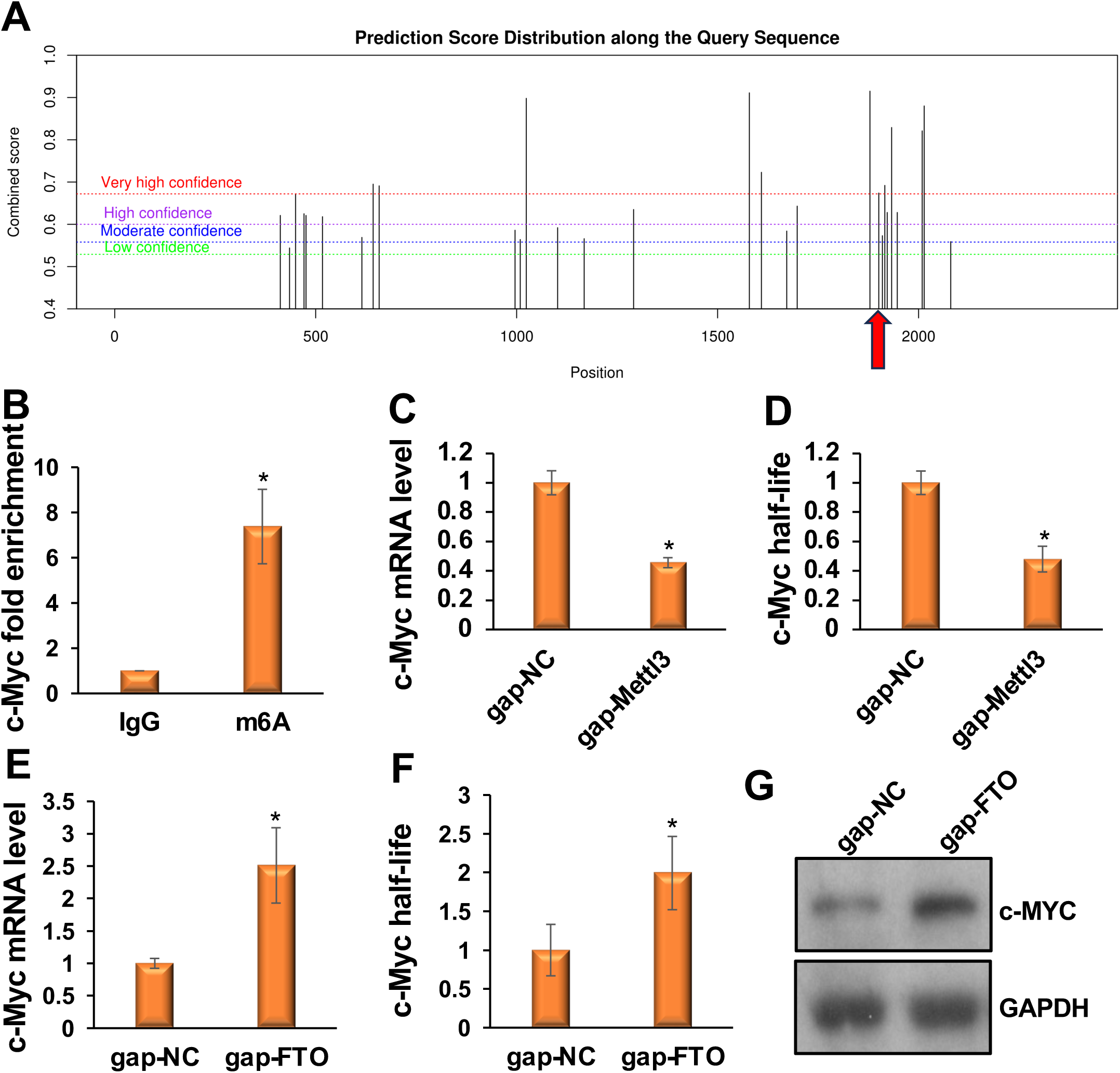
FTO promotes myogenesis by demethylating m^6^A-modified *c-Myc* and regulating its expression. (**A**) The SRAMP algorithm detected a cluster of highly confident m^6^A sites near the stop codon of the *c-Myc* transcript. A red arrow marks the position of the stop codon. (**B**) m^6^A-RIP qPCR shows that *c-Myc* mRNAs are highly enriched when pulled down with the m^6^A antibody compared with the IgG control. (**C, D**) Knockdown of *Mettl3* decreases *c-Myc* transcript levels and half-lives. (**E, F**) Knockdown of *Fto* increases *c-Myc* transcript levels and half-lives. (**G**) Knockdown of *Fto* increases c-MYC protein levels. Values are expressed as mean ± SD of biological triplicates. *P < 0.001.

To determine whether *c-Myc* is indeed m^6^A-modified, we performed RNA immunoprecipitation with an m^6^A antibody followed by qRT-PCR with *c-Myc*-specific primers (m^6^A-RIP qPCR), following a standard protocol^43^. We immunoprecipitated ribosome-depleted RNAs from myoblasts with either an m^6^A antibody or an IgG control, then performed qRT-PCR with c-Myc-specific primers. *c-Myc* mRNAs were highly enriched in the m^6^A immunoprecipitated samples compared with the IgG control (**Fig. 4B**). Next, we assessed the effect of m^6^A levels on *c-Myc* transcripts and their half-lives. We knocked down *Mettl3* in myoblasts and measured *c-Myc* levels. Mettl3 knockdown decreased *c-Myc* levels (**Fig. 4C**). We also determined the half-life of *c-Myc* in *Mettl3* knockdown myoblasts using a standard protocol and found it was shortened in these cells (**Fig. 4D**). These findings confirm that METTL3 installs the m^6^A epitranscriptomic mark on *c-Myc*. To further determine whether *c-Myc* is regulated by FTO, we knocked down *Fto* in primary myoblasts. *Fto* knockdown increased *c-Myc* transcripts, their half-life, and protein levels (**Fig. 4E-G**). Taken together, these studies demonstrate that FTO promotes myogenesis by regulating m^6^A-modified *c-Myc*.

### IGF2BP2 reader protein recognizes m^6^A-modified c-Myc and stabilizes it

Specific reader proteins recognize m^6^A-modified RNAs and regulate their fate by modulating transport, splicing, decay, and translation^15,18–23^. A newly identified reader protein, insulin-like growth factor 2 mRNA-binding protein 2 (IGF2BP2), has been shown to stabilize m^6^A-modified RNAs^18^. We hypothesize that FTO levels increase during myoblast differentiation, thereby decreasing m^6^A levels on c-Myc. This reduction prevents the IGF2BP2 reader protein from binding *c-Myc*, thereby destabilizing *c-Myc* and promoting differentiation. To test this, we first verified whether IGF2BP2 binds to the *c-Myc* transcript in myoblasts. We performed RNA immunoprecipitation using ribosome-depleted RNAs from myoblasts with either an m^6^A antibody or an IgG control. We then performed qRT-PCR on these samples using *c-Myc*-specific primers. *c-Myc* mRNA was highly enriched in the m^6^A immunoprecipitated samples compared with the IgG control (**Fig. 5A**). We knocked down *Igf2bp2* in myoblasts and measured *c-Myc* transcript levels. *Igf2bp2* knockdown decreased *c-Myc* transcript levels (**Fig. 5B**). We also determined the half-life of *c-Myc* in *Igf2bp2* knockdown myoblasts using a standard protocol and found it was shortened in these cells (**Fig. 5C**). As expected, c-MYC protein levels also decreased in the *Igf2bp2* knockdown myoblasts (**Fig. 5D**). These findings confirm that the IGF2BP2 reader protein binds to m^6^A-modified *c-Myc* and stabilizes it in undifferentiated myoblasts. During myoblast differentiation, m^6^A levels on *c-Myc* decrease due to increased FTO levels, which prevents IGF2BP2 from binding to *c-Myc*, thereby destabilizing *c-Myc* and promoting differentiation.

**Figure 5:**
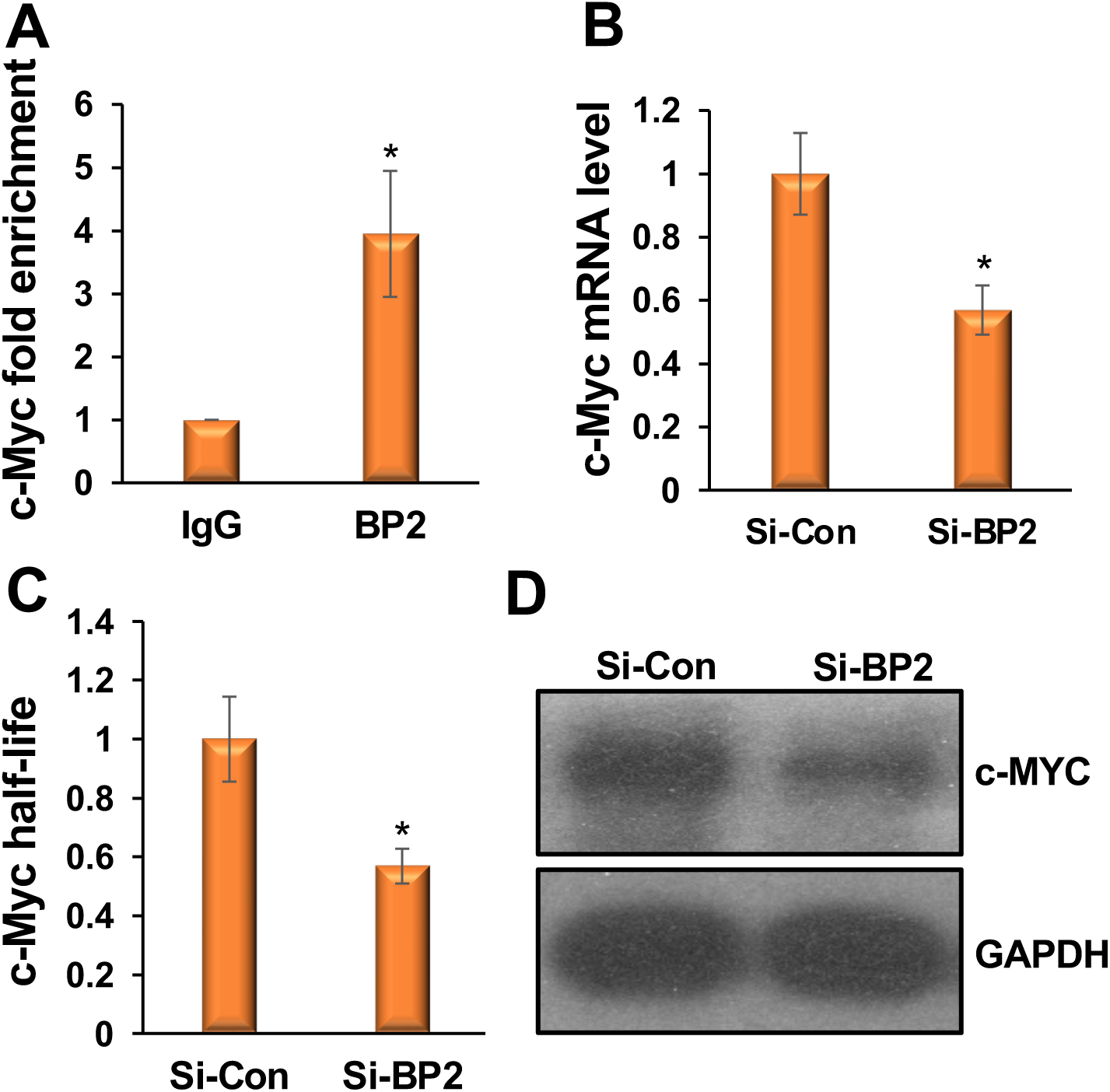
The IGF2BP2 reader protein binds m^6^A-modified *c-Myc* and stabilizes it. (**A**) RNA immunoprecipitation qPCR shows that *c-Myc* mRNA was highly enriched when pulled down with the IGF2BP2 (BP2) antibody compared with the IgG control. (**B**) Knockdown of *Igf2bp2* decreases *c-Myc* mRNA levels. (**C**) An RNA stability assay shows that *Igf2bp2* knockdown after actinomycin D treatment reduces *c-Myc* mRNA half-lives. (**D**) Knockdown of *Igf2bp2* decreases c-MYC protein levels. Values are expressed as mean ± SD of biological triplicates. (A) *P < 0.005. (B, C) *P < 0.001.

### FTO is required for skeletal muscle regeneration

We used a well-established mouse model to study skeletal muscle regeneration. In this model, we induce injury by injecting cardiotoxin (CTX) into the tibialis anterior (TA) muscle of adult mice. The injury initially causes muscle degeneration, followed by spontaneous regeneration, allowing us to study the process of muscle regeneration (**Fig. 1D**). As previously shown, *Fto* levels sharply decrease during the first 1 to 3 days after CTX injury (the degeneration phase) and then increase as new muscle fibers are formed (the regeneration phase) (**Fig. 1E**). This regulation of *Fto* during both the degeneration and regeneration phases suggests that it plays a vital role in the critical processes of muscle regeneration.

We investigated the role of FTO in skeletal muscle regeneration using *Fto* flox (*Fto^f/f^*) mice. As is well known, new muscle fibers are generated from MuSCs. FTO is expressed in MuSCs. Therefore, to investigate its role in muscle regeneration, we knocked out *Fto* in a MuSC-specific manner. We generated an *Fto* conditional knockout (cKO) mouse line (*Pax7^CreERT2^*; *Fto^f/f^*) by crossing *Pax7^CreERT2^* with *Fto^f/f^*. We administered intraperitoneal injections of tamoxifen to *Fto* cKO mice and littermate controls (*Pax7^CreERT2^* and *Fto^f/f^*) at approximately 9 weeks of age, at 100 mg/kg body weight for five consecutive days^44,45^. Subsequently, we injected 10 μM CTX in PBS (100 μL) into the TA muscle belly of these mice, following our established protocol^46–48^.

By day 5, regeneration was in its early stage, and new myofibers were beginning to form. Therefore, we examined the effects of *Fto* cKO on days 7 and 14 post-injury. We harvested the TA muscles and performed H&E staining on fresh-frozen muscle sections using our established protocol. In the control group, muscle regeneration occurred normally, and by day 14, morphology was restored. However, central nuclei, a hallmark of regenerating muscle, were present (**Fig. 6A**). In contrast, muscle regeneration was impaired in *Fto* cKO mice (**Fig. 6A**). We also observed an increased number of nuclei, indicating an influx of inflammatory cells alongside other cell types(**Fig. 6A**). Additionally, smaller myofibers were present, suggesting that regeneration was compromised (**Fig. 6A**).

**Figure 6:**
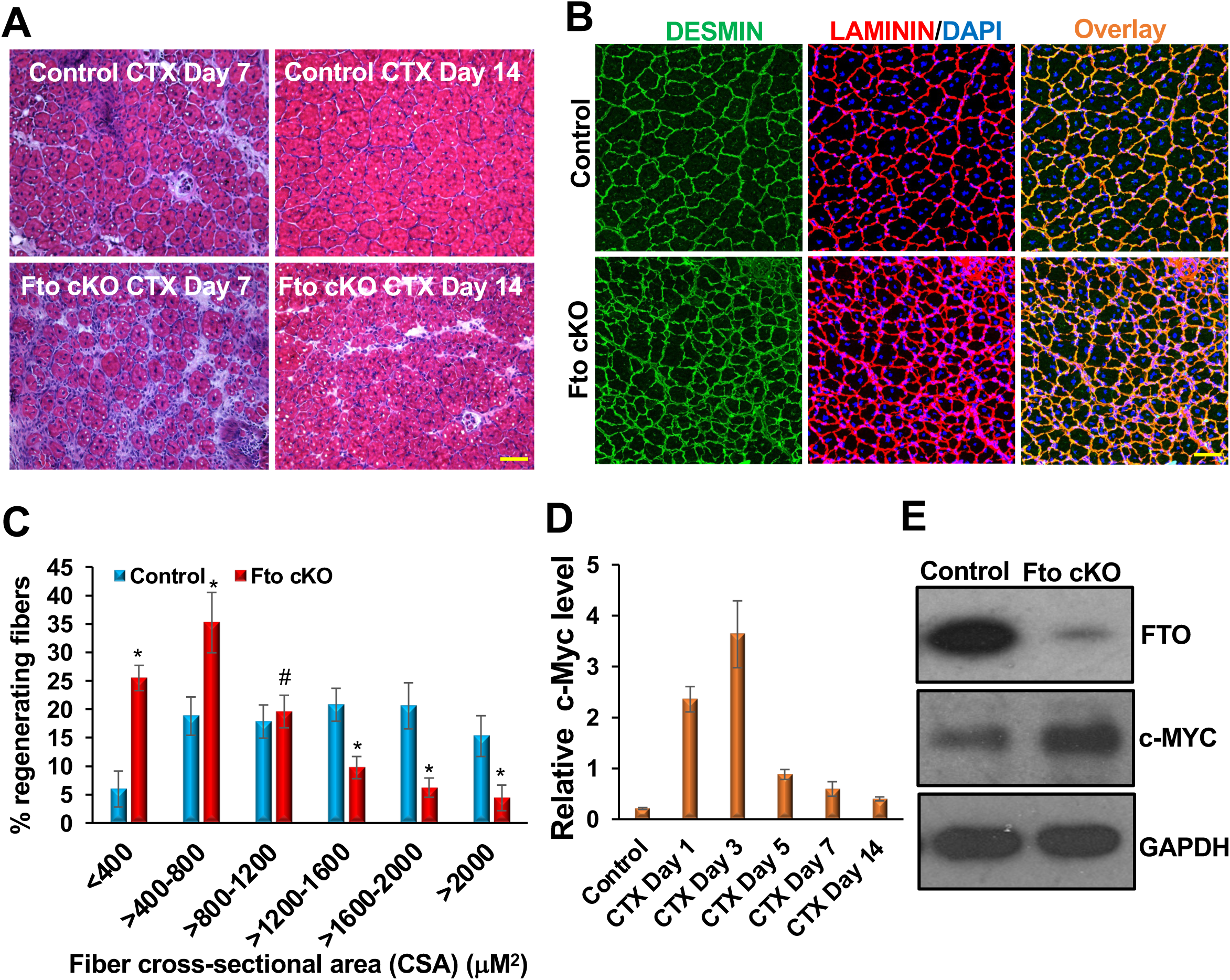
FTO is required for skeletal muscle regeneration. (**A)** Representative H&E images show impaired muscle regeneration in *Fto* cKO mice on days 7 and 14 after CTX injury. Smaller muscle fibers and an increased number of nuclei, indicating an influx of inflammatory cells, were observed. (**B**) Representative DESMIN and LAMININ staining of muscle sections on day 14 after CTX injury. Muscle fibers were significantly smaller, the spaces between fibers were less well defined, and the sections were filled with numerous nuclei in *Fto* cKO mice. (**C**) Cross-sectional areas (CSA) of regenerating fibers on day 14 were measured using Fiji. There was a significant increase in the number of smaller CSAs, while the number of larger CSAs decreased significantly in *Fto* cKO samples. (**D**) *c-Myc* is gradually upregulated in TA muscles on days 1–3 post-injury and downregulated on days 5–14 post-injury. (**E**) c-MYC is derepressed in *Fto* cKO TA muscles 14 days after injury. Scale bar: 50 μM. Mean ± SD from samples of five mice. *P < 0.001; ^#^P = non-significant.

We further performed immunostaining of muscle sections from day 14 post-injury for DESMIN and LAMININ. DESMIN is an intermediate filament protein highly expressed in newly formed muscle fibers, whereas LAMININ labels the boundaries of these fibers. We observed intense expression of both DESMIN and LAMININ in all samples (**Fig. 6B**). However, the morphology of muscle sections from *Fto* cKO samples appeared abnormal. The muscle fibers were significantly smaller than those in control samples (**Fig. 6B**). Additionally, the spaces between fibers were less well defined and filled with numerous nuclei (**Fig. 6B**). We used Fiji software to measure the cross-sectional areas (CSAs) of the myofibers. We observed a dramatic increase in the number of smaller CSAs, while the number of larger CSAs decreased significantly in the *Fto* cKO samples (**Fig. 6C**), further indicating impaired regeneration. These results suggest that FTO is essential for proper skeletal muscle regeneration *in vivo*.

Furthermore, we examined the levels of FTO target *c-Myc* during regeneration and observed that its expression pattern was inversely related to *Fto* (**Figure 1E**; **Fig. 6D**). Specifically, *c-Myc* levels increased during the initial 1 to 3 days post-injury and then steadily declined from days 5 to 14 after the injury (**Fig. 6D**). Additionally, c-MYC protein levels were derepressed in the *Fto* cKO TA muscle samples (**Fig. 6E**). These observations indicate that FTO plays a critical role in regulating *c-Myc* during the regeneration process. These studies collectively suggest that the novel FTO/c-Myc axis is crucial for normal skeletal muscle regeneration *in vivo*.

## Discussions

Myogenesis occurs in a coordinated manner and requires strict regulation of gene expression to maintain normal skeletal muscle function. Abnormalities in the gene regulation of critical myogenic pathways can lead to numerous life-threatening muscle degenerative diseases. Over the past several decades, research on myogenesis has primarily focused on how myogenic transcription factors and signaling molecules regulate this process^49–51^. However, the mechanisms that regulate these molecules remain largely unknown, suggesting that many essential regulators of myogenic processes have yet to be identified. For example, the roles of epigenetics^52^ and non-coding RNAs^46,53–56^ in skeletal muscle biology are now well established. More recently, post-transcriptional (epitranscriptomic) modifications have emerged as a vital aspect of gene regulation in various biological processes and pathological conditions^57,58^. To address this knowledge gap, we have investigated the role of an epitranscriptomic regulator in myoblast differentiation and skeletal muscle regeneration.

In this study, we elucidate the crucial role of the m^6^A RNA demethylase FTO and its underlying molecular mechanisms in myoblast differentiation and skeletal muscle regeneration in mice. Our findings reveal that FTO is significantly upregulated during both the differentiation and regeneration phases of skeletal muscle. Exogenous FTO expression enhances myoblast differentiation, whereas FTO inhibition suppresses it. Depleting FTO in MuSCs of the TA muscles in mice impairs muscle regeneration. FTO promotes these processes by targeting and regulating m^6^A-modified *c-Myc*, a repressor of myogenesis, with the help of the IGF2BP2 reader protein, which stabilizes m^6^A-modified *c-Myc* in undifferentiated myoblasts. Increased FTO levels during differentiation reduce m^6^A modifications on *c-Myc*, thereby preventing its stabilization by the IGF2BP2 reader protein and promoting differentiation. Overall, our findings emphasize the vital role of the novel FTO/c-Myc/IGF2BP2 axis in regulating skeletal muscle differentiation and regeneration.

Thus far, most studies have demonstrated the role of the m^6^A RNA modification writer METTL3 in myogenesis. METTL3 regulates MuSC/myoblast state transitions^25^, *Myod* mRNA maintenance in proliferating myoblasts^26^, myoblast differentiation^25,27,28^, regeneration^27^, skeletal muscle maintenance and growth^29^, and fat intake in skeletal muscles^30^. However, studies implicating the m^6^A demethylase FTO in myogenesis are limited. Moreover, these studies are largely limited to *in vitro* myogenesis^24,32^. Our study is crucial because it reveals the role of FTO in myogenesis using mouse primary myoblasts and a well-established mouse model of regeneration.

MuSCs are the source of new muscle fibers. MuSCs typically remain quiescent in resting muscle. However, these quiescent cells are activated following injury, giving rise to myoblasts. These myoblasts then differentiate and fuse with existing myofibers to repair the injured myofiber or form new myofibers. FTO is expressed in MuSCs, persists through the myoblast stage, and is upregulated during myoblast differentiation. Here, we depleted FTO levels during the MuSC stage and revealed its role in regeneration. We have also shown that FTO is required for normal myoblast differentiation, suggesting that it is required at the myoblast stage for normal muscle regeneration. We will address this vital question in our future studies.

More significantly, we have established *c-Myc*, an essential regulator of myogenesis, as a direct target of FTO in these processes. c-MYC is a transcription factor that binds E-boxes^42^, underscoring its critical role in regulating myogenic processes. As is well known, the myogenic master regulator, MYOD, regulates the myogenic program by binding E-boxes in critical myogenic factors^59^. Consistent with this, recent studies have shown that *c-Myc* regulates myogenesis by targeting prominent myogenic microRNAs and non-coding RNAs^42^, further suggesting that *c-Myc* converges on a crucial node and may serve as a primary direct target of FTO in myogenesis.

Mechanistically, we have revealed that the new m^6^A reader protein IGF2BP2 regulates the stability of m^6^A-modified *c-Myc* during myogenesis. Most m^6^A reader proteins, such as YTHDF2^19^, degrade their target molecules. Here, we showed that IGF2BP2 stabilizes *c-Myc* in undifferentiated myoblasts when FTO expression is low. During differentiation, FTO levels increase, preventing *c-Myc* from being m^6^A-modified and thereby preventing its stabilization by the IGF2BP2 reader protein, thereby promoting differentiation. Overall, our findings emphasize the vital role of the novel FTO/c-Myc/IGF2BP2 axis in regulating skeletal muscle differentiation and regeneration. This complex and critical role of FTO in regeneration will help in understanding devastating muscle degenerative diseases.

## Materials and methods

### Cell culture

We obtained the C2C12 mouse myoblast cell line from the American Type Culture Collection (ATCC). We cultured these cells at subconfluent density in growth medium (GM) composed of DMEM from ATCC, 10% heat-inactivated fetal bovine serum (FBS) from ATCC, and 1X Antibiotic-Antimycotic from Thermo Scientific. For differentiation into myocytes or myotubes, we used the differentiation medium (DM) Field ^53^. The DM was composed of DMEM with 2% heat-inactivated horse serum from Hyclone and 1X Antibiotic-Antimycotic from Thermo Scientific. Additionally, mouse primary myoblasts were isolated from C57BL/6J mice following a standard protocol previously detailed^60^. We subsequently differentiated these primary myoblasts using the procedures outlined earlier^61^.

### Isolation of total RNA and performance of qRT–PCR

Total RNA was isolated using Trizol reagent (Invitrogen) or the RNeasy mini kit (Qiagen) following the manufacturer’s instructions. cDNA synthesis was performed using the iScript cDNA Synthesis Kit (Bio-Rad) following the manufacturer’s instructions. qRT-PCR was performed using the SYBR Green PCR master mix (Bio-Rad) on a Bio-Rad thermal cycler. The specific primers used were: Myogenin (*Myog*) (Forward: AGCGCAGGCTC AAGAAAGTGAATG; Reverse: CTGTAGGCGCTCAAT GTACTGGAT), Myosin Heavy Chain (*Mhc*) (forward: TCC AAACCGTCTCTGCACTGTT; Reverse: AGCGTACAA AGTGTGGGTGTGT). We used *Rsp13* (Forward: TGACGACGTGAA GGAACAG ATTT; Reverse: ATTTCCAGTCACAAAACG GACCT) or *Gapdh* (Forward: ATGACATCAAGAAG GTG GTGAAGC; Reverse: GAAGAGTGGGAGTTGCTGTTG AAG) primer pairs as housekeeping genes to normalize the values of the genes of interest.

### Western blotting and antibodies

We harvested the cells, washed them with 1X PBS, lysed them in NP-40 lysis buffer (50 mM Tris-HCl, 150 mM NaCl, 0.1% NP-40, 5 mM EDTA, 10% glycerol) supplemented with a protease inhibitor cocktail (Sigma). We separated proteins by SDS-PAGE, transferred them to nitrocellulose, and immunoblotted with the following antibodies: anti-MYOG (1:500; Santa Cruz), anti-MHC (1:3,000; Sigma), anti-c-MYC (1:1,000; Santa Cruz), and anti-GAPDH (1:10,000; Sigma).

### Skeletal muscle regeneration studies

We purchased C57BL/6J (Stock #000664), *Fto* flox (Stock #027830), and Pax7CreERT2 (Stock #017763) from Jackson Laboratories. We performed our mouse work in our animal facility under an approved Institutional Animal Care and Use Committee (IACUC) protocol. We generated an *Fto* conditional knockout (cKO) mouse line (*Pax7^CreERT2^*; *Fto^f/f^*) by crossing *Pax7^CreERT2^* with *Fto^f/f^*. We administered intraperitoneal injections of tamoxifen to *Fto* cKO mice and littermate controls (*Pax7^CreERT2^* and *Fto^f/f^*) at approximately 9 weeks of age, at 100 mg/kg body weight for five consecutive days^44,45^. Subsequently, we studied muscle regeneration using a well-established mouse model generated by injecting cardiotoxin (CTX) from Naja nigricollis (EMD Millipore), following the procedure described earlier^46–48^. Briefly, we injected the TA muscles of about 10-week-old mice with 100 μL of 10 μM CTX. The high volume of injected material, pressure, and post-injection massage spread the material throughout the TA muscle compartment. We analyzed only the middle two-thirds of the TA muscle closest to the injection site. We euthanized the mice by CO2 and then sacrificed them by cervical dislocation before harvesting the muscle samples at different time intervals.

### Immunocytochemistry and immunohistochemistry

Immunocytochemistry was performed using our standard protocol, as described earlier [14, 27]. Briefly, cells were cultured on sterile glass coverslips and fixed with 2% formaldehyde in PBS for 15 min. Next, cells were permeabilized with 0.2% Triton X-100 and 1% normal goat serum (NGS) in ice-cold PBS for 5 min, then blocked with 1% NGS in PBS twice for 15 min each. Subsequently, cells were incubated with primary antibodies in 1% NGS for one hour. We used the MYOG antibody from Santa Cruz Biotechnology at 1:50 and the MHC antibody from Sigma at 1:400. After these steps, cells were washed twice with 1X PBS and incubated with a FITC-conjugated anti-mouse IgG secondary antibody for 1 hour. We obtained this antibody from DAKO Cytomation and used a 1:500 dilution. Finally, cells were washed twice, and nuclei were counterstained with DAPI (Vector Laboratories). Coverslips were mounted on glass slides, and images were captured using a Leica fluorescence microscope. Immunostaining of TA muscle tissue sections was performed using the previously optimized protocol. We used primary antibodies rat anti-LAMININ (1:100; Millipore) and mouse anti-DESMIN (1:200; Dako). Secondary antibodies Alexa Fluor 594 goat anti-mouse IgG1 (1:400; Invitrogen) and Alexa Fluor 488 goat anti-rabbit IgG (1:400; Invitrogen) were used. Images were captured using a Zeiss LSM-710 confocal microscope. H&E staining was performed according to the standard protocol, and bright-field images were acquired using a Leica microscope.

### Statistical analyses

We presented our data as the mean ± standard deviation of three or more biological replicates. A two-tailed Student’s *t*-test was employed to determine P-values.

## Acknowledgments

This work was supported by an NIH grant (1R21AR080936-01A1) to BKD.

## Author contributions

BKD conceived the project; PD and BKD designed the study, performed the experiments, analyzed the data, and prepared the manuscript.

## Supplemental Tables

**Supplemental Figure 1:**
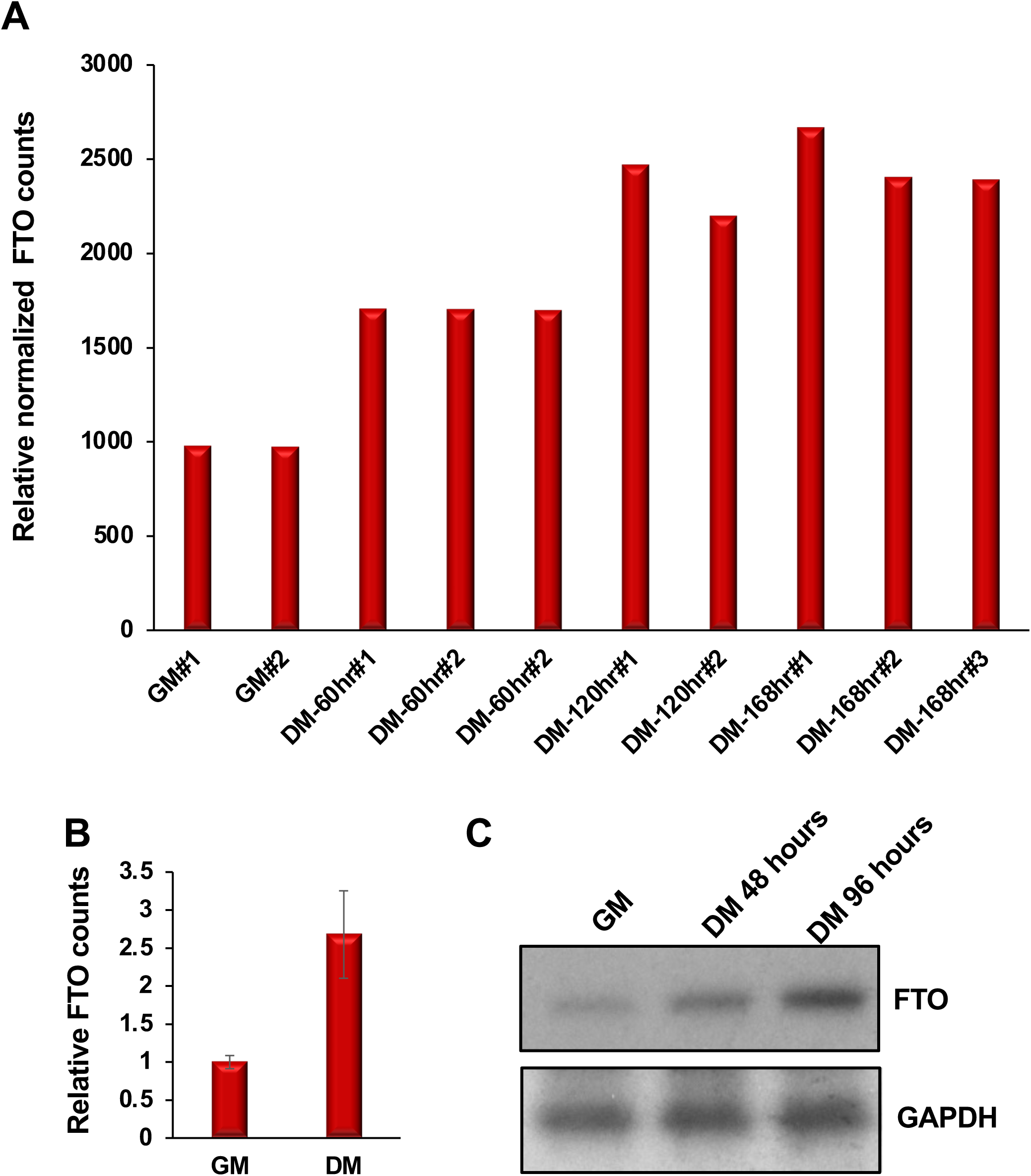
FTO is upregulated during C2C12 myoblast differentiation. (**A**) Normalized RNA-seq counts from undifferentiated C2C12 myoblasts (GM) and myoblasts differentiated in differentiation medium (DM) for 60, 120, and 168 hours are shown. (**B**) qRT-PCR analyses show that *Fto* is upregulated during C2C12 myoblast differentiation. *Fto* levels were first normalized to *Gapdh*, and fold changes were calculated relative to undifferentiated myoblasts (GM). DM indicates culture in differentiation medium for 96 hours. (**C**) FTO protein levels, as shown by Western blotting, also increase during mouse C2C12 myoblast differentiation. GAPDH served as a loading control.

**Supplemental Figure 2:**
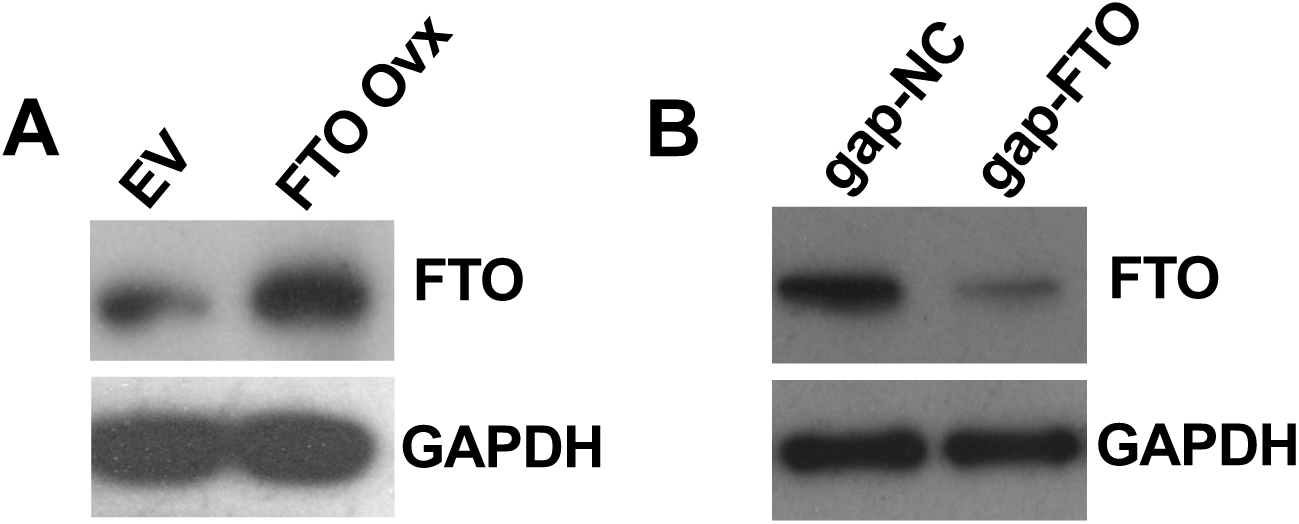
(**A**) Western blotting showing that myoblasts transduced with a retroviral vector expressing FTO increase FTO protein level. (**B**) Western blot analysis showing that myoblasts treated with gap-FTO decrease FTO protein levels.

## Supplemental Figures

**Supplemental Table 1:** Ectopic expression of FTO increases the number of MYOG- and MHC-positive cells.

| Treatments | MYOG-positive cell number |  |  | MHC-positive cell number |  |  |
| --- | --- | --- | --- | --- | --- | --- |
|  | Total Cells<br>(12 fields) | Cells per field |  | Total Cells<br>(12 fields) | Cells per field |  |
|  |  | Mean | SD |  | Mean | SD |
| EV | 467 | 38.91 | 5.68 | 395 | 32.91 | 4.42 |
| FTO Ovx | 1012 | 84.33 | 8.26 | 828 | 69.00 | 5.86 |

**Supplemental Table 2:** Knockdown of *Fto* decreases the number of MYOG- and MHC-positive cells.

Dey\_Supplemental Table 2
| Treatments | MYOG-positive cell number |  |  | MHC-positive cell number |  |  |
| --- | --- | --- | --- | --- | --- | --- |
|  | Total Cells<br>(12 fields) | Cells per field |  | Total Cells<br>(12 fields) | Cells per field |  |
|  |  | Mean | SD |  | Mean | SD |
| gap-NC | 550 | 45.83 | 8.35 | 536 | 44.47 | 5.91 |
| gap-FTO | 286 | 23.83 | 5.07 | 250 | 20.83 | 3.15 |

